# Quinolone resistance genes *qnr*, *aac(6′)-Ib-cr*, *oqxAB*, and *qepA* in environmental *Escherichia coli*: insights into their genetic contexts from comparative genomics

**DOI:** 10.1101/2024.04.24.590863

**Authors:** Ryota Gomi, Fumie Adachi

## Abstract

Previous studies have reported the occurrence of transferable quinolone resistance determinants in environmental *Escherichia coli*. However, little is known about their vectors and genetic contexts. To gain insights into these genetic characteristics, we analyzed the complete genomes of 53 environmental *E. coli* isolates containing one or more transferable quinolone resistance determinants, including 20 sequenced in this study and 33 sourced from RefSeq. The studied genomes carried the following transferable quinolone resistance determinants alone or in combination: *aac(6′)-Ib-cr*, *oqxAB*, *qepA1*, *qnrA1*, *qnrB4*, *qnrB7*, *qnrB19*, *qnrD1*, *qnrS1*, and *qnrS2*, with *qnrS1* being predominant. These resistance genes were detected on plasmids of diverse replicon types; however, *aac(6′)-Ib-cr*, *qnrS1*, and *qnrS2* were also detected on the chromosome. The genetic contexts surrounding these genes included not only those found in clinical isolates but also novel contexts, such as *qnrD1* embedded within a composite transposon-like structure bounded by Tn*3*-derived inverted-repeat miniature elements (TIMEs). This study provides deep insights into mobile genetic elements associated with transferable quinolone resistance determinants, highlighting the importance of genomic surveillance of antimicrobial resistant bacteria in the environment.

## 1. INTRODUCTION

*Escherichia coli* is a commensal member of the gut microbiota of humans and animals but can also cause intestinal and extraintestinal infections.^1^ Another concern besides pathogenicity is the rise of antimicrobial resistance (AMR) in *E. coli*.^2^ This is due to the accumulation of AMR determinants, including acquired AMR genes and chromosomal point mutations.^3^

Quinolones are antibiotics that are widely used for treatment of infections caused by a variety of bacteria, including *E. coli*.^4^ Quinolone resistance in *E. coli* is mainly attributed to mutations in the quinolone resistance-determining regions (QRDRs) of the topoisomerase genes *gyrA* and *parC*.^5,6^ Although these chromosomal mutations are generally not horizontally transferred, there are quinolone resistance determinants that can be transmitted between bacteria, including target protection (*qnr*), antibiotic efflux (*qepA* and *oqxAB*), and antibiotic modification (*aac(6′)-Ib-cr*).^7^ These transferable resistance determinants are called plasmid-mediated quinolone resistance (PMQR) genes or transferable mechanisms of quinolone resistance (TMQR) determinants (a recent review recommended the use of TMQR because these genes are sometimes present on the chromosome, so we use the term TMQR in this paper).^7^ It is known that the levels of resistance conferred by TMQR are relatively low; however, TMQR can play an important role in the development of quinolone resistance because (i) the effect of TMQR on quinolone minimum inhibitory concentrations (MICs) is additive to that conferred by other transferable or chromosome-mediated quinolone resistance determinants, and (ii) TMQR can facilitate the selection of mutants with higher-level quinolone resistance.^7,8^

Information on the vectors and genetic contexts of AMR genes is important for multiple reasons: for example, (i) it allows identification of frequently associated/linked resistance genes and thus can provide information on co-resistance; (ii) it facilitates better understanding of the evolution of multiresistance regions and how resistance genes spread.^9,10^ Previous studies reported the occurrence of TMQR determinants in environmental *E. coli*, and some studies also reported the sequences of replicons carrying TMQR determinants.^11,12^ However, in-depth analysis of the genetic contexts of TMQR determinants has not been performed in most cases, and whether the environmental isolates carry TMQR determinants in the contexts related to those in clinical isolates or in previously unreported contexts is largely unknown. Here, we analyzed the complete genomes of environmental *E. coli* isolates, sequenced in this study or obtained from the NCBI reference sequence (RefSeq) database, with the aim to acquire genomic insights into TMQR, including their vectors and genetic contexts.

## 2. MATERIALS AND METHODS

### 2.1. *E. coli* isolates

Twenty-four environmental *E. coli* isolates carrying TMQR determinants isolated in Japan were characterized in this study. These 24 isolates included 19 isolates from our previous collections: (i) three isolates from river water samples collected in 2010 (1002W03, 1008S13, and 1008S19),^13^ (ii) 10 isolates from river water samples collected between 2011–2013 (KFu015, KMi012, KMi014, KMi029, KKa014, KFu021, KOr024, KTa005, KTa007, KTa009),^14^ and (iii) six extended-spectrum β-lactamase (ESBL)-producing *E. coli* isolates obtained from municipal wastewater (JSWP006, JSWP021) and hospital wastewater (JKHS004, JKHS006, JKHS016, and JKHS019) in 2015.^15^ Five isolates were obtained in the present study: three isolates were screened from *E. coli* isolated from municipal wastewater using CHROMagar ECC (Kanto Chemical Co., Tokyo, Japan) supplemented with cefotaxime (1 mg/L) between 2019–2020 (19M19, 19A14, and 20A24), and two isolates were screened from *E. coli* isolated from river water using CHROMagar ECC supplemented with ciprofloxacin (0.03 mg/L) between 2021–2022 (21F61 and 22F46). For the newly obtained isolates, the presence of TMQR genes was confirmed by PCR using primers described previously,^13,16,17^ with a slight modification (5′-GTGAAGTCGATCAGTCAGTG-3′ was used instead of oqxAFw).

### 2.2. Antimicrobial susceptibility testing

Antimicrobial susceptibility testing of the 24 isolates was performed by microdilution using Dry Plate Eiken (Eiken, Tokyo, Japan) according to CLSI specifications (see **Table S1** for antimicrobials used). The MIC of ciprofloxacin was also determined using the Etest (bioMérieux, Marcy-l’Étoile, France). The results were evaluated according to CLSI criteria^18^ and EUCAST epidemiological cutoff (ECOFF) values (https://mic.eucast.org/search/).

### 2.3. Genome sequencing and assembly

The genomes of 24 isolates were sequenced using a combination of short-read (Illumina) and long-read (Oxford Nanopore Technologies) sequencing. Hybrid assembly of Illumina short reads and ONT long reads was performed using both a long-read-first approach and a short-read-first approach. The details of genome sequencing and assembly are described in the Supplementary Materials and methods. Of the 24 isolates, the genomes of 20 isolates could be completed, while the genomes of four isolates (KMi014, KTa007, KTa009, JKHS019) could not be completed despite repeated long-read sequencing. The 20 completed genomes were further analyzed to elucidate the genetic contexts of TMQR determinants.

### 2.4. Retrieving complete *E. coli* genomes with TMQR determinants from the NCBI database

RefSeq *E. coli* genomes with an assembly level of “complete genome” (n = 2815) were downloaded using the ncbi-genome-download tool (v0.3.1, https://github.com/kblin/ncbi-genome-download) in July 2023. Metadata information, including strain name and isolation source, was extracted from the downloaded files, and genomes determined to be of environmental origin (i.e., surface water, sediment, wastewater, and soil) (n = 203) were retained. AMR genes were detected using ABRicate (v1.0.1, https://github.com/tseemann/abricate) with the NCBI database.^19^ This identified 48 genomes with at least one TMQR determinant. For each of the 48 genomes, we searched PubMed and Google Scholar for the associated publication. This identified publications for 33 genomes, which were further analyzed in this study. Detailed information on these 33 genomes is summarized in **Table S2**.

### 2.5. Genomic analysis

The *E. coli* genomes sequenced in this study and those retrieved from RefSeq were subjected to the following genomic analysis. AMR genes were detected using ResFinder 4.1^20^ and ABRicate (v1.0.1, https://github.com/tseemann/abricate) with the NCBI database.^19^ Multilocus sequence typing (MLST) was performed using mlst (v2.19.0, https://github.com/tseemann/mlst). Plasmid replicons were detected using PlasmidFinder 2.1, and plasmids were typed using pMLST 2.0.^21^ Genomes were annotated using the RAST server,^22^ ISfinder,^23^ and the blastn program (https://blast.ncbi.nlm.nih.gov/Blast.cgi) with the core nucleotide (core_nt) database (last accessed in August 2024).

### 2.6. Data availability

The complete genomes and sequence reads obtained in the present study have been deposited in GenBank and the NCBI SRA under BioProject PRJNA1078256 (also see **Table S1** for the accession number of each genome).

## 3. RESULTS AND DISCUSSION

### 3.1. Basic characteristics of *E. coli* isolates sequenced in this study

Twenty-four *E. coli* isolates with TMQR determinants were sequenced in the present study. These *E. coli* isolates belonged to 20 different STs, including clinically important lineages such as ST131 (**Table S1**). These isolates carried the following TMQR determinants alone or in combination: *qnrS1* (n = 12), *qnrD1* (n = 6), *aac(6′)-Ib-cr* (n = 4), *qnrB7* (n = 3), *qnrS2* (n = 3), *qepA1* (n = 1), *qnrB19* (n = 1), and *oqxAB* (n = 1) (**Table S1**).

The isolates were non-susceptible to a wide variety of antibiotics (**Table S1**). Nonsusceptibility rates for quinolones were 29.2% for nalidixic acid, 37.5% for ciprofloxacin (evaluated by Etest), and 37.5% for levofloxacin. All isolates were non-susceptible to nalidixic acid, ciprofloxacin, and levofloxacin when they carried a QRDR mutation(s). Isolates with TMQR determinants but without QRDR mutations were susceptible to nalidixic acid but either susceptible or non-susceptible to ciprofloxacin and levofloxacin. This is consistent with previous studies reporting that isolates with TMQR determinants but without QRDR mutations sometimes show an unusual phenotype of nalidixic acid susceptibility and ciprofloxacin resistance.^7^ Isolates with TMQR determinants but without QRDR mutations were all classified as non-wild type to ciprofloxacin and levofloxacin according to the ECOFF criteria, indicating that interpreting results with ECOFF values can increase the sensitivity for detection of TMQR determinants.

### 3.2. Basic characteristics of RefSeq *E. coli* genomes with TMQR determinants

Thirty-three RefSeq *E. coli* genomes with TMQR determinants were retrieved, which were associated with 12 publications. These were from wastewater (n = 18) or environmental water (n = 15), isolated between 2012 and 2022, and from various geographical locations, with Switzerland (n = 14 from two publications) and Japan (n = 10 from three publications) being predominant (**Table S2**). In total, 24 different STs were detected among the genomes. These genomes carried the following transferable quinolone resistance determinants alone or in combination: *qnrS1* (n = 21), *aac(6′)-Ib-cr* (n = 9), *qnrB4* (n = 3), *oqxAB* (n = 3), *qnrS2* (n = 2), *qnrA1* (n = 1), and *qnrB19* (n = 1) (**Table S2**). Although we extracted RefSeq genomes associated with publications, the genetic contexts of TMQR genes were investigated only for four genomes in four publications (**Table S2**). Even for these genomes, the contexts were often insufficiently or incorrectly annotated. We thus performed in-depth analysis of the genetic contexts in the RefSeq genomes together with our genomes as below.

### 3.3. Genetic contexts of *qnrA1*

A RefSeq genome analyzed in this study carried *qnrA1* on an IncA/C plasmid, p009_A. IS*CR1* is usually present upstream of *qnrA1*, whereas downstream structures of *qnrA1* are known to be divergent.^24^ In p009_A, *qnrA1* was embedded in a structure related to class 1 integrons In36 and In37 (**Figure 1**).^25^ A structure related to another class 1 integron, In4873, was detected upstream of *qnrA1*.^26^ The association of In36/In37-like structure and In4873-like structure seems to be rare: blastn analysis found only one closely related structure in an IncHI2/IncHI2A plasmid, pME-1a, from a clinical *Enterobacter hormaechei* isolate from the USA (**Figure 1**).^27^ It is assumed that these closely related structures derive from the same ancestral structure, which was (i) separately introduced into an IncA/C plasmid and an IncHI2/IncHI2A or (ii) generated in or introduced into one plasmid, which was then mobilized to the other.

**Figure 1.**
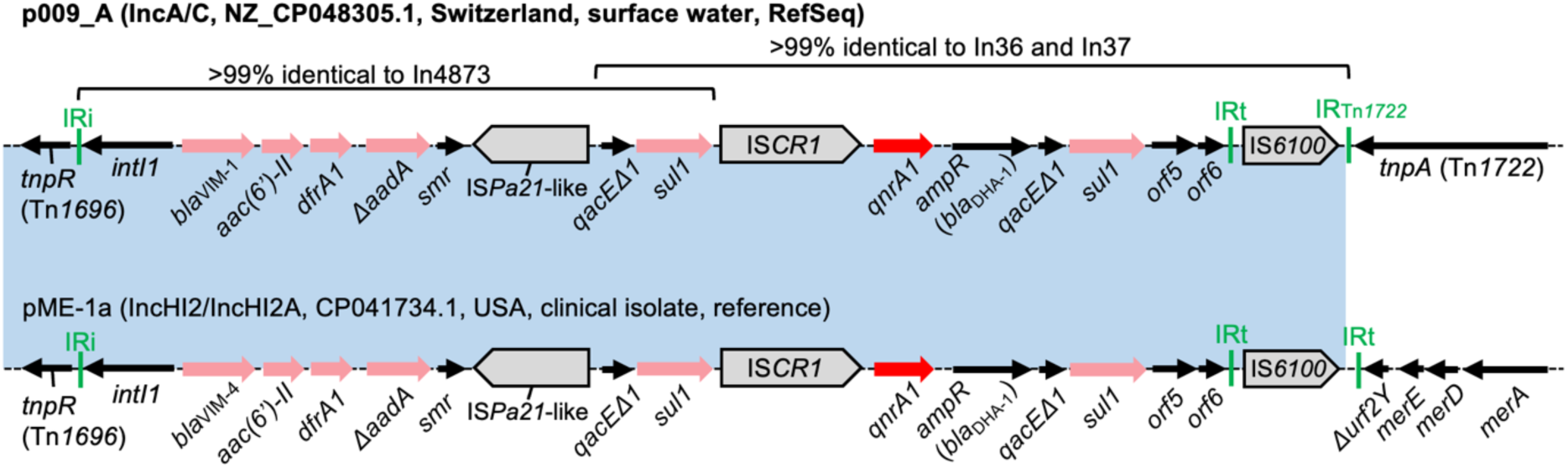
Genetic contexts of *qnrA1*. Red arrows indicate TMQR determinants, pink arrows indicate other AMR genes, gray pointed boxes indicate insertion sequences (ISs), and black arrows indicate other genes. Diagrams are not exactly to scale (this also applies to the other figures). Inc type, nucleotide accession number, country of origin, isolation source, and sequence note are shown in parentheses next to the plasmid name. The context detected in pME-1a is shown for reference purposes. pME-1a carries *bla*_VIM-4_ instead of *bla*_VIM-1_. The blue shaded regions indicate nucleotide sequences with >99% identity. IR: inverted repeat. IRi: inverted repeat at *intI1* end. IRt: inverted repeat at *tni* end.

### 3.4. Genetic contexts of *qnrB4*

Three RefSeq genomes carried *qnrB4* on different IncF plasmids. *qnrB4* is known to be globally distributed and often associated with the AmpC type β-lactamase gene *bla*_DHA-1_.^24^ In all three plasmids, *qnrB4* is linked to *bla*_DHA-1_ and also to other AMR genes (**Figure 2a**). The context from *sul1* to the truncated class 1 integron was detected in multiple clinical isolates (e.g., CP058590), but the broader context seen in p142_A-OXA181, which links *qnrB4* and *qnrS1*, was not detected in any other public genomes by blastn analysis. The effect on susceptibility to quinolones by carrying multiple *qnr* genes is not completely understood but seems to be minor, because the products of two genes can compete for binding to gyrase.^24^

**Figure 2.**
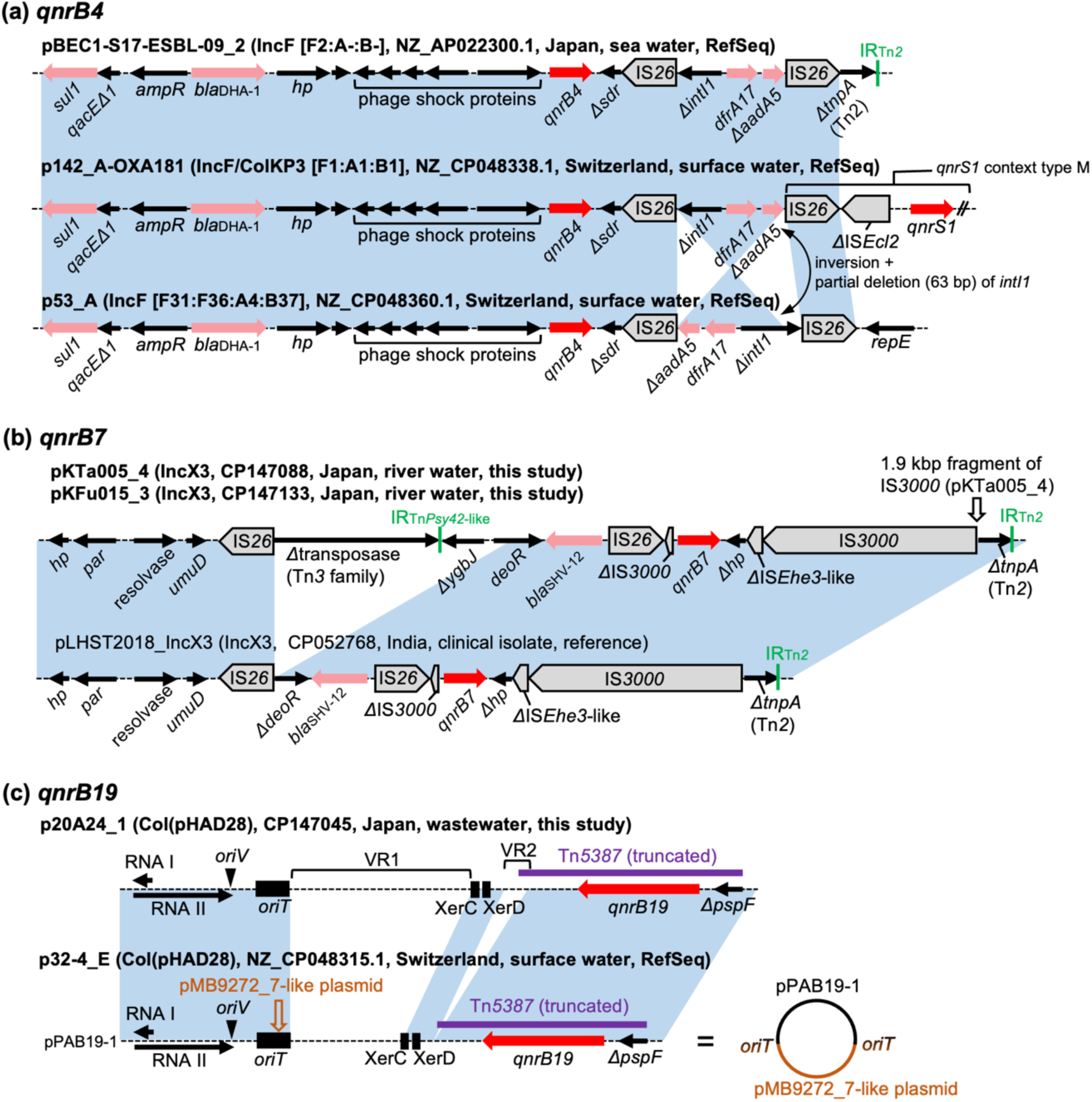
Genetic contexts of *qnrB* genes. Genes and elements are shown as in Figure 1. IncF replicon sequence types are indicated in square brackets. Nucleotide sequences with >99% identity are indicated by the blue shaded regions. (a) Genetic contexts of *qnrB4*. In p53_A, the *ΔintI1*-*dfrA17*-*ΔaadA5* structure is inverted and the *ΔintI1* sequence is 63-bp longer compared with the other two plasmids. (b) Genetic contexts of *qnrB7*. The contexts are almost identical in pKTa005_4 and pKFu015_3 except that a 1.9 kbp fragment of IS*3000* is inserted next to another IS*3000* in pKTa005_4. The context in pLHST2018_IncX3 is shown for reference purposes. (c) Genetic contexts of *qnrB19*. Panel (c) is enlarged compared with panels (a) and (b), and describes the whole plasmid structures. p20A24_1 has undergone modifications in variable regions called VR1 and VR2 in comparison to the common plasmid pPAB19-1.^29^ A circular structure of p32-4_E is also shown to describe the relationship to pPAB19-1 and the pMB9272_7-like plasmid. The putative ColE1-like *ori* (RNAI, RNAII, and *oriV*), *oriT*, and Xer sites (sites used for converting plasmid multimers into monomers) are indicated. The *oriT* sequence in the pMB9272_7-like plasmid was predicted by comparison with well-characterized plasmids such as ColE1. hp: hypothetical protein.

### 3.5. Genetic contexts of *qnrB7*

Two isolates sequenced to closure in the present study carried *qnrB7*. *qnrB7* was previously reported to be associated with the ESBL gene *bla*_SHV-12_.^24^ However, reports of complete plasmid (or chromosome) sequences carrying *qnrB7* are scarce; blastn analysis identified only four complete plasmid sequences carrying *qnrB7*. One of them, the IncX3 plasmid pLHST2018_IncX3 harbored by a clinical *Salmonella enterica* isolate from India, carried both *qnrB7* and *bla*_SHV-12_.^28^ In our isolates, *qnrB7* was also carried by IncX3 plasmids and associated with *bla*_SHV-12_ (**Figure 2b**). The genetic contexts in our isolates and pLHST2018_IncX3 were similar, but a region next to *bla*_SHV-12_ was deleted in pLHST2018_IncX3. Detection of closely related AMR plasmids in environmental and clinical bacteria from different countries highlights the importance of global One Health surveillance to track the spread of AMR genes.

### 3.6. Genetic contexts of *qnrB19*

Two genomes, one sequenced by us and the other from RefSeq, carried *qnrB19*. *qnrB19* is usually mobilized by transposition units (TPU) mediated by IS*Ecp1*, such as Tn*2012* and Tn*5387*, or small ColE1-like plasmids containing fragments of these transposons.^29^ The most common *qnrB19*-carrying ColE1-like plasmid is a 2699 bp plasmid named pPAB19-1 (or pSGI15/pECY6-7/pMK100), with others being minor variants of this plasmid.^29^ The two *qnrB19*-positive genomes carried this gene on small ColE1-like plasmids, p20A24_1 and p32-4_E. p20A24_1 is related to the common plasmid pPAB19-1, with modifications in two regions (**Figure 2c**). p32-4_E was identified as a cointegrate comprising pPAB19-1 and a 1718 bp plasmid almost identical (single nucleotide difference) to pMB9272_7 (CP103528), with putative *oriT* regions being the cointegration points. No close hits were found for this plasmid by blastn analysis (all hits were below 80% query coverage). pMB9272_7 carries only two genes encoding hypothetical proteins, and no plasmid replicon was identified by PlasmidFinder. Cointegrates comprising ColE1-like plasmids and large plasmids were previously shown to play an important role in the spread of AMR.^30^ The example of p32-4_E indicates cointegrates of ColE1-like plasmids and other small plasmids may also contribute to the spread of AMR.

### 3.7. Genetic contexts of *qnrD1*

Three genomes sequenced to closure in the present study carried *qnrD1*. *qnrD1* was first described on a 4270 bp plasmid, p2007057, in a clinical *Salmonella enterica* strain isolated in China.^31^ *qnrD1* has since been detected on multiple plasmids; however, blastn analysis identified that *qnrD1* is rare in *Enterobacteriaceae* and mostly detected on ∼2.7 kbp plasmids in *Morganellaceae*. Two of the three genomes carried *qnrD1* on a plasmid identical to p2007057 (**Figure 3**). The remaining genome carried *qnrD1* on a 6657 bp plasmid, named pKFu015_4, with no close blastn hits (all hits were below 50% query coverage). The *qnrD1* region, including *qnrD1* and ORF2, in pKFu015_4 is identical to ∼2.7 kbp *Morganellaceae* plasmids such as pRS12-11 (KF364953) and is ∼95% identical to the corresponding region in p2007057 (and thus pKMi029_6 and pKTa005_7). Analysis of sequences surrounding the *qnrD1* region revealed 209 bp directly oriented repeats upstream and downstream of the region. This repeat carries imperfect 33 bp inverted repeats (IRs) closely related to some Tn*3*-family transposons (e.g., Tn*Ec4*), though it does not encode a transposase. The blastn analysis revealed that this repeat is identical to a Tn*3*-derived inverted-repeat miniature element (TIME), named TIME_IS*101*_.^32^ Pairs of TIMEs are known to be involved in mobilization of intervening sequences when the corresponding Tn*3* family transposases are provided *in trans*, and these composite transposon-like structures were previously named TIME-COMP.^32,33^ Thus, the TIME-COMP structure, TIME_IS*101*_-*qnrD1*-ORF2-TIME_IS*101*_, seems to have been mobilized as a single unit into a ColE1-like plasmid, generating pKFu015_4. The presence of 5 bp direct repeats (AGCTA) surrounding this structure supports this idea. Moreover, we also found a ColE1-like plasmid without insertion of this TIME-COMP structure among the genomes sequenced in this study (pKMi029_5, CP147114), further supporting this. *qnrD1* was previously suggested to be mobilized as a mobile insertion cassette (mic) element (a nonautonomous element bracketed by two IRs),^34^ but this observation indicates another mode of *qnrD1* mobilization, namely TIME-COMP.

**Figure 3.**
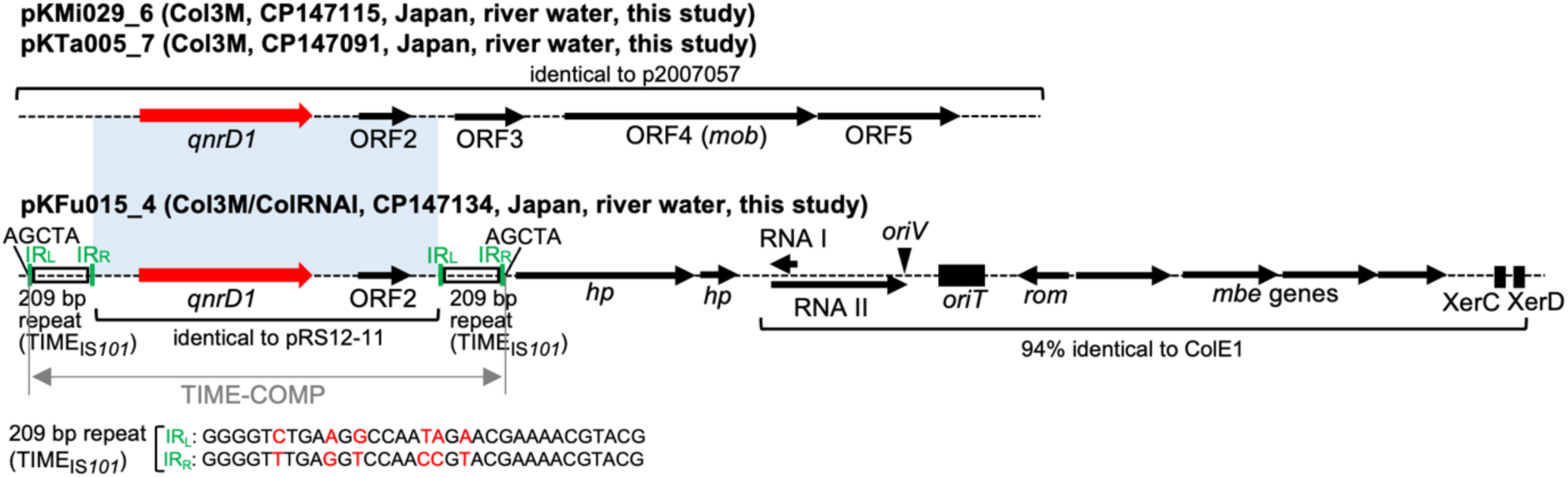
Structures of plasmids carrying *qnrD1*. Genes and elements are shown as in earlier figures. The light blue shaded regions indicate nucleotide sequences with ∼95% identity. pKFu015_4 carries regions corresponding to the *ori*, *bom* (basis of mobility), *rom* (RNA one inhibition modulator), *mob*, and *cer* (ColE1 resolution sequence) regions in plasmid ColE1.^51^ IR_L_: inverted repeat left. IR_R_: inverted repeat right. IR_L_ and IR_R_ of TIME*_IS101_* are shown at the bottom of the figure, and different nucleotides are marked in red.

### 3.8. Genetic contexts of *qnrS1*

*qnrS1* was found to be the most prevalent TMQR gene both in our isolates (n = 12) and in the RefSeq genomes (n = 21) and was detected on plasmids of diverse replicon types or on the chromosome (**Table S1** and **Table S2**). Although earlier studies showed low *qnrS1* prevalence,^7^ some recent studies identified *qnrS1* as the most prevalent TMQR gene in the One Health context.^35,36^ The genetic contexts of *qnrS1* were largely classified into 13 types (types A to M in **Figure 4**). Types B, E, and I are found only in the genomes analyzed in this study and no other hits were found by blastn analysis. On the other hand, the other genetic contexts were found in at least one public genome, and some were described in previous studies (e.g., types A, F and G previously described in clinical or animal isolates).^37–41^ *qnrS1* is frequently linked to other resistance genes in the contexts, including an ESBL gene *bla*_CTX-M-15_ in types B–D. The potential evolutionary pathways of these context types are described in **Figure S1**. Briefly, IS*26*-mediated rearrangements and IS*Ecp1*-mediated transposition seem to have played major roles in diversification of the contexts and linkage of *qnrS1* to other AMR genes.

**Figure 4.**
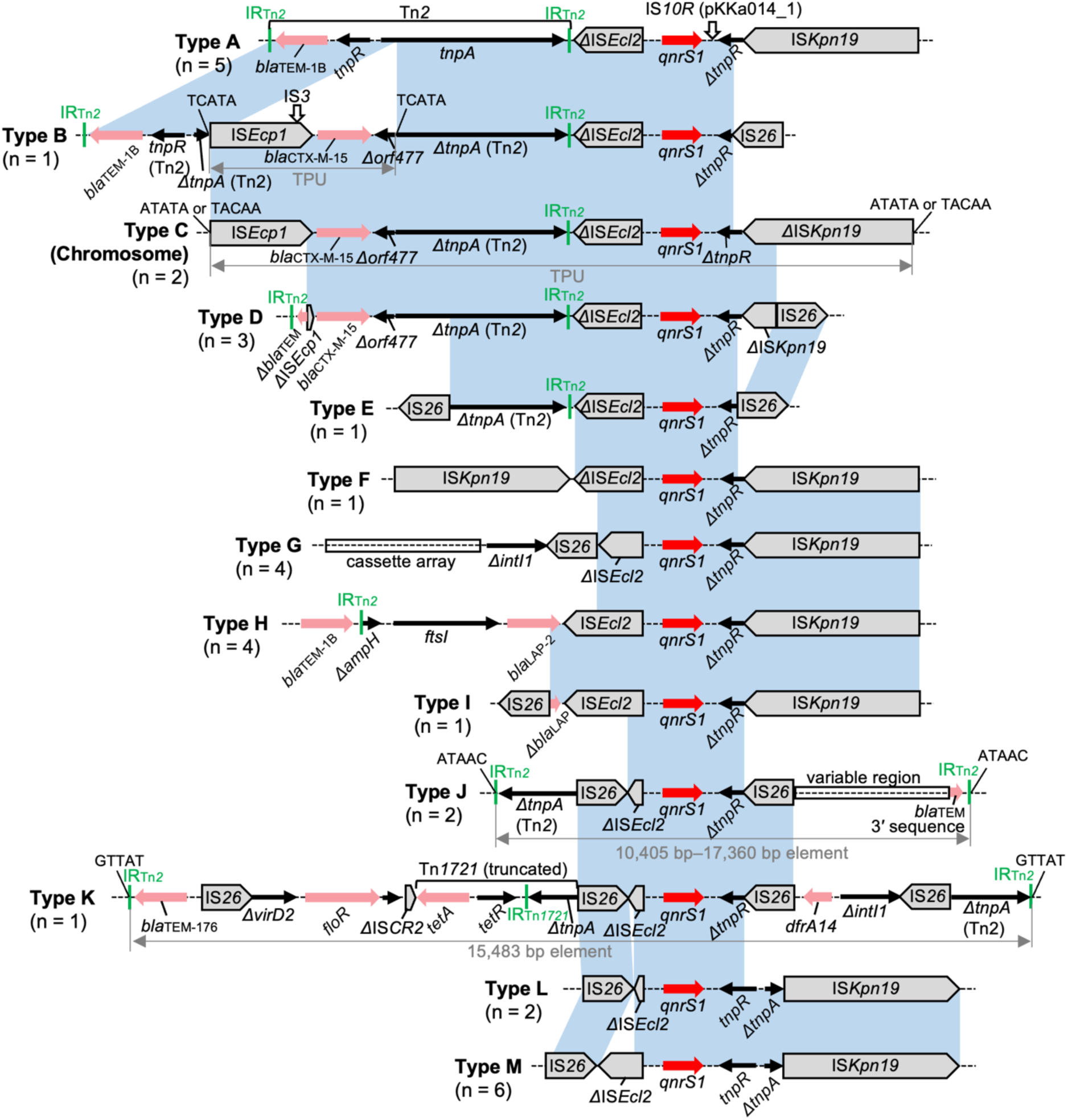
Genetic contexts of *qnrS1*. Genes and elements are shown as in earlier figures. The blue shaded regions indicate nucleotide sequences with >99% identity. Note that for simplicity, only *qnrS1* and directly flanking regions without complex rearrangements (such as inversions) are shaded in blue (e.g., the left IS*Kpn19* in the type F context shares >99% nucleotide identity with IS*Kpn19* in the type G context, but this region is not shaded). The number of genomes carrying each context type is shown in parentheses. See **Table S1** and **Table S2** for information on genomes carrying each context type, and **Figure S1** for possible evolutionary pathways of these contexts.

### 3.9. Genetic contexts of *qnrS2*

Two genomes sequenced to closure in this study and two RefSeq genomes carried *qnrS2*. *qnrS2* was previously reported to be part of a mic element bracketed by 22 bp imperfect IRs.^42^ In all four genomes, the mic elements containing *qnrS2* were truncated (**Figure 5**). Two RefSeq genomes carried *qnrS2* in the same genetic context on IncX1 plasmids. The *qnrS2* region in these plasmids was followed by the *fosA3* genetic context type K,^43^ linking *qnrS2* to *fosA3*, *Δbla*_TEM_, and *bla*_CTX-M-55_. This linkage was not detected in any other public genomes by blastn analysis. Two ST2179 genomes sequenced in this study carried *qnrS2* on the chromosome. In 19M19, the *qnrS2* region was duplicated. The blastn search identified five *E. coli* genomes carrying *qnrS2* on the chromosome, three of which belong to ST2179. The genetic contexts of *qnrS2* in these three genomes were closely related to that in 19M19, but the *qnrS2* region was not duplicated (**Figure 5**). It is assumed that a resistance region containing *qnrS2* was inserted into an ancestral ST2179 genome, which was followed by IS*26*-mediated rearrangements, yielding the variation of the contexts seen in **Figure 5**.

**Figure 5.**
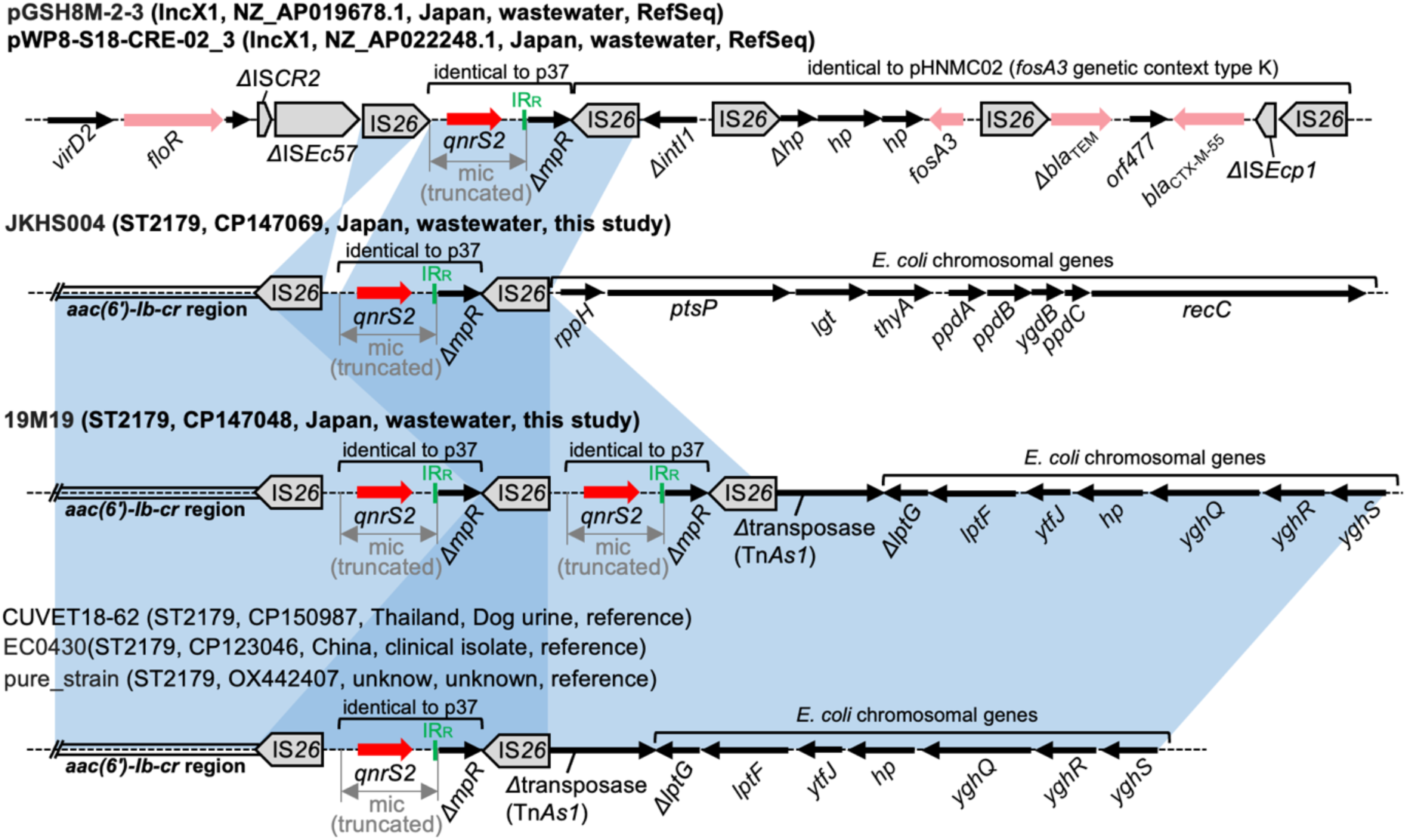
Genetic contexts of *qnrS2*. Genes and elements are shown as in earlier figures. JKHS004 and 19M19 carry *qnrS2* on the chromosome, and chromosomal STs are indicated instead of plasmid types. The blue shaded regions indicate nucleotide sequences with >99% identity. The shared context detected in CUVET18-62, EC0430, and pure_strain is shown for reference purposes. The chromosomal IS*26*-*qnrS2*-*ΔmpR*-IS*26* structure in ST2179 genomes is different from that in the IncX1 plasmids, in that the IS*26* sequences are directly-oriented and a ∼200 bp fragment is present between the truncated mic and the left IS*26*.

### 3.10. Genetic contexts of *aac(6′)-Ib-cr*

Three genomes sequenced to closure in this study and nine RefSeq genomes carried *aac(6′)-Ib-cr*. *aac(6′)-Ib-cr* was detected on the chromosome (n = 6) or on IncF plasmids (n = 6) (**Figure 6**). The isolates with chromosomal *aac(6′)-Ib-cr* belonged to five different STs (ST10, ST410, ST648, ST1266, and ST2179). A chromosomal location for *aac(6′)-Ib-cr* was also reported previously in clinical ST131 and ST648 isolates from Spain.^44^ These examples indicate chromosomal integration of *aac(6′)-Ib-cr* has occurred multiple times in different STs.

**Figure 6.**
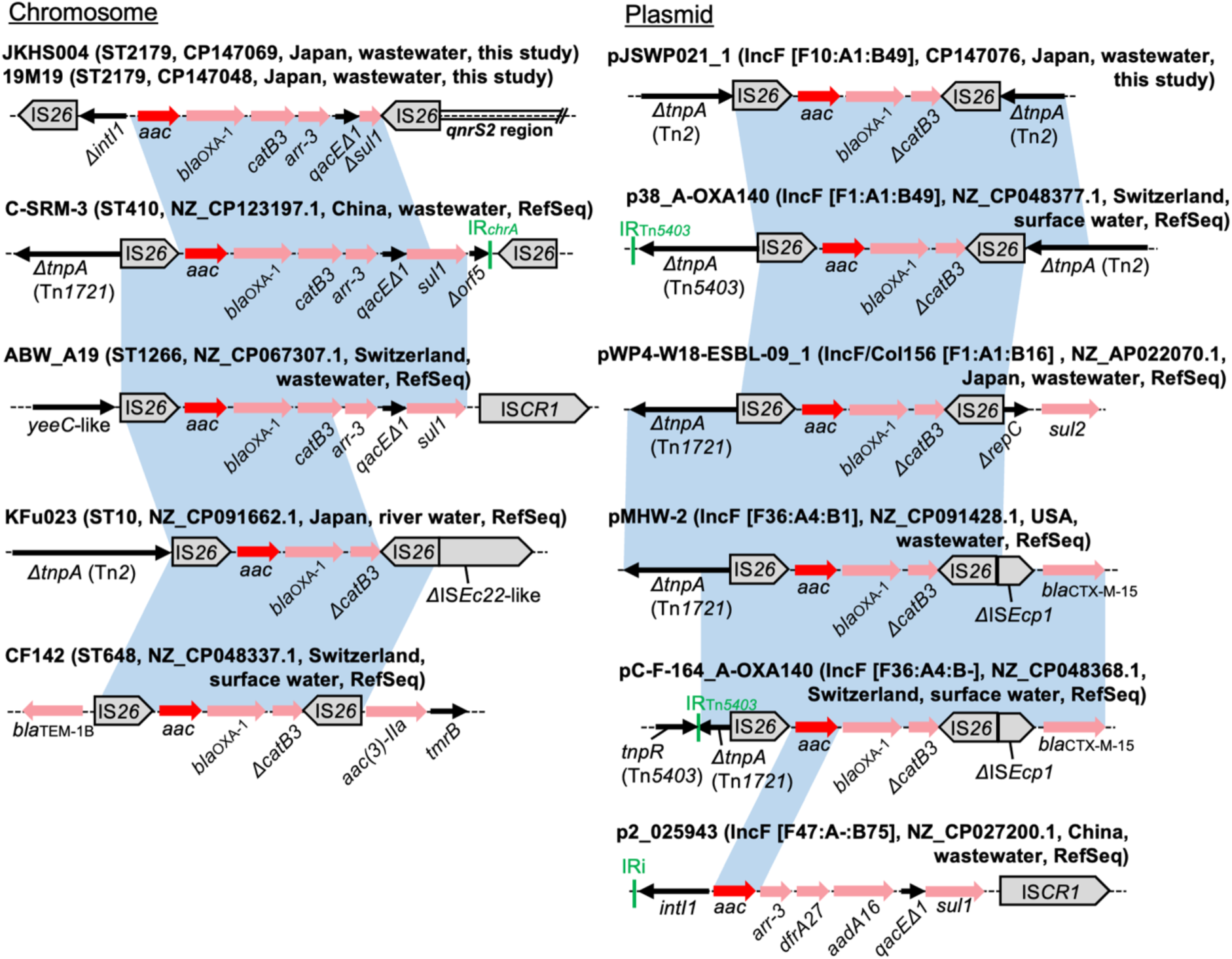
Genetic contexts of *aac(6′)-Ib-cr*. The contexts are shown separately for chromosomes and plasmids. *aac(6′)-Ib-cr* is abbreviated as *aac* in the figure. Genes and elements are shown as in earlier figures. The blue shaded regions indicate nucleotide sequences with >99% identity. Note that for simplicity, only *aac(6′)-Ib-cr* and flanking regions are shaded in blue (e.g., IS*26* sequences in JKHS004 and 19M19 share >99% nucleotide identity to IS*26* sequences in C-SRM-3, but these regions are not shaded). In all replicons except p2_025943, either or both of the 5′-conserved segment and the 3′-conserved segment of class 1 integrons containing *aac(6′)-Ib-cr* are interrupted by IS*26*. These truncated integrons are likely to have originated from the class 1 integron *intI1*-*aac(6′)-Ib-cr*-*bla*_OXA-1_-*catB3*-*arr-3*-*qacEΔ1*-*sul1*.

*aac(6′)-Ib-cr* has been identified in a gene cassette as a segment of class 1 integrons.^45^ The IS*26*-*aac(6′)-Ib-cr*-*bla*_OXA-1_-*ΔcatB3*-IS*26* structure was prevalent and detected in seven genomes (58%), while the upstream and downstream sequences were divergent (**Figure 6**). This structure is also common in public genomes (more than 1000 hits by blastn). However, the IS*26* sequences are inversely oriented and thus this structure is neither a transposon nor a pseudo-compound transposon.^46^ The prevalence of this structure among various plasmids/chromosomes can partially be explained by the presence of other mobile genetic elements surrounding it, which might have aided the movement of multiresistance regions containing this structure.

### 3.11. Genetic contexts of *oqxAB*

Mobile *oqxAB* genes are commonly found within an IS*26*-*oqxA*-*oqxB*-*oqxR*-IS*26* pseudo-compound transposon, named Tn*6010*.^47^ All four *oqxAB* genes identified in the present study are situated within Tn*6010* and carried by plasmids of different Inc types (**Figure 7**). The sequences upstream of Tn*6010* (in reference to the orientation of *oqxAB* genes) in three RefSeq plasmids share some common regions, which were also detected upstream of Tn*6010* in multiple public genomes by blastn. On the other hand, the upstream sequence of Tn*6010* was replaced by a truncated Tn*1721* and a truncated Tn*2* in pJKHS004_1, linking *tetA* and a truncated *bla*_TEM_ to *oqxAB*. No genomes were found by blastn analysis to carry this multiresistance region.

**Figure 7.**
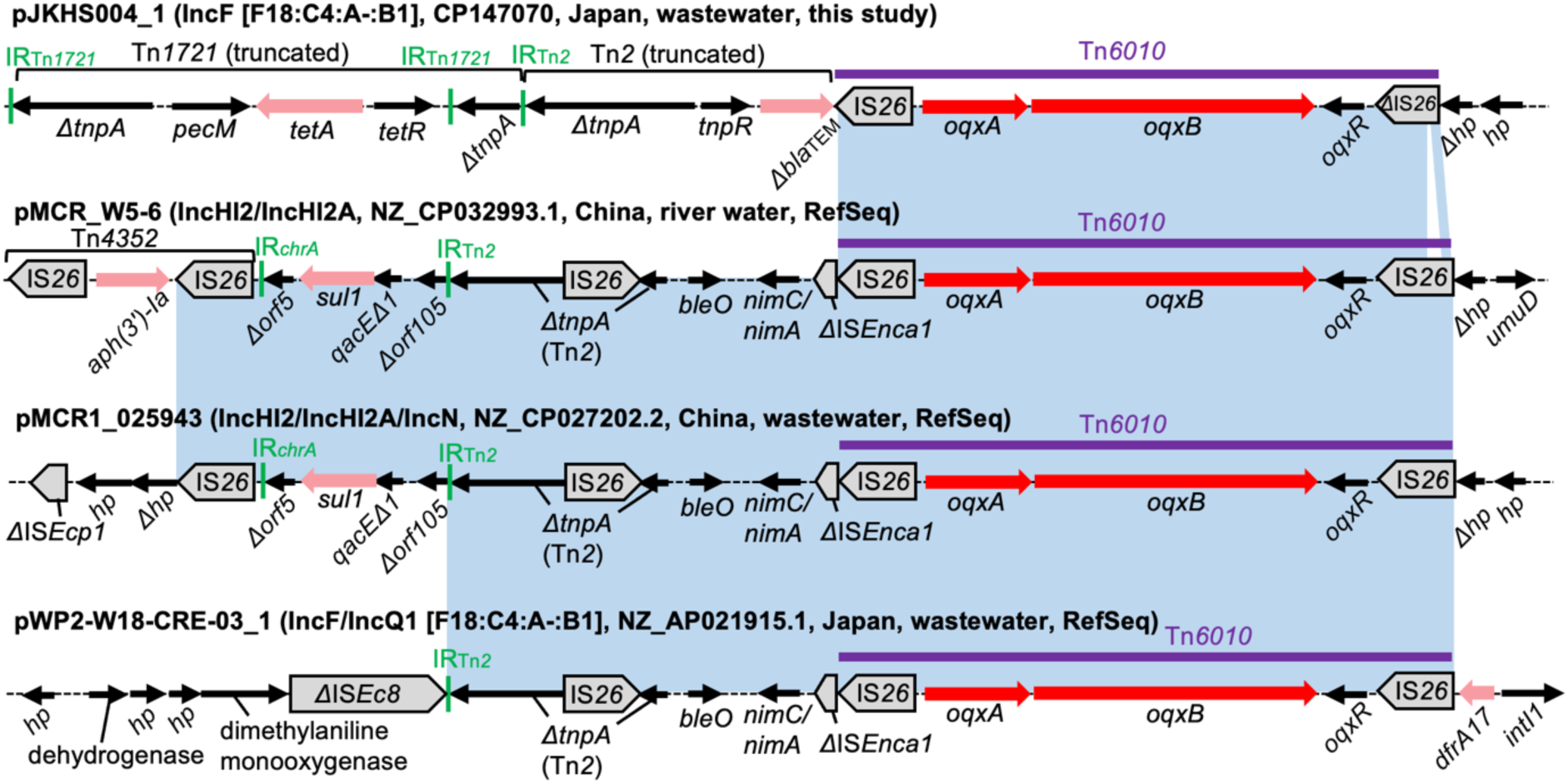
Genetic contexts of *oqxAB*. Genes and elements are shown as in earlier figures. The blue shaded regions indicate nucleotide sequences with >99% identity. Note that for simplicity, only Tn*6010* and flanking regions are shaded in blue (e.g., the left IS*26* in Tn*4352* shares >99% nucleotide identity to other IS*26* sequences, but these regions are not shaded). Tn*6010* in pJKHS004_1 has an internally deleted copy of IS*26*.

### 3.12. Genetic contexts of *qepA1*

*qepA* genes are typically associated with IS*CR3* and embedded within complex integrons.^48^ One isolate sequenced in this study carried *qepA1*, and the gene was located between a truncated *intI1* and IS*CR3* on an IncF plasmid (**Figure 8**). Other AMR genes, *dfrB4*, *sul1*, and *mph(A)*, were found to be linked to *qepA1* in this context. The blastn analysis of the entire resistance region bounded by IS*26* revealed closely related sequences in three *E. coli* plasmids, one from a clinical isolate in the USA and two from clinical isolates in Myanmar,^49,50^ indicating intercontinental occurrence of this multiresistance region and highlighting the need for a One Health approach to AMR surveillance.

**Figure 8.**
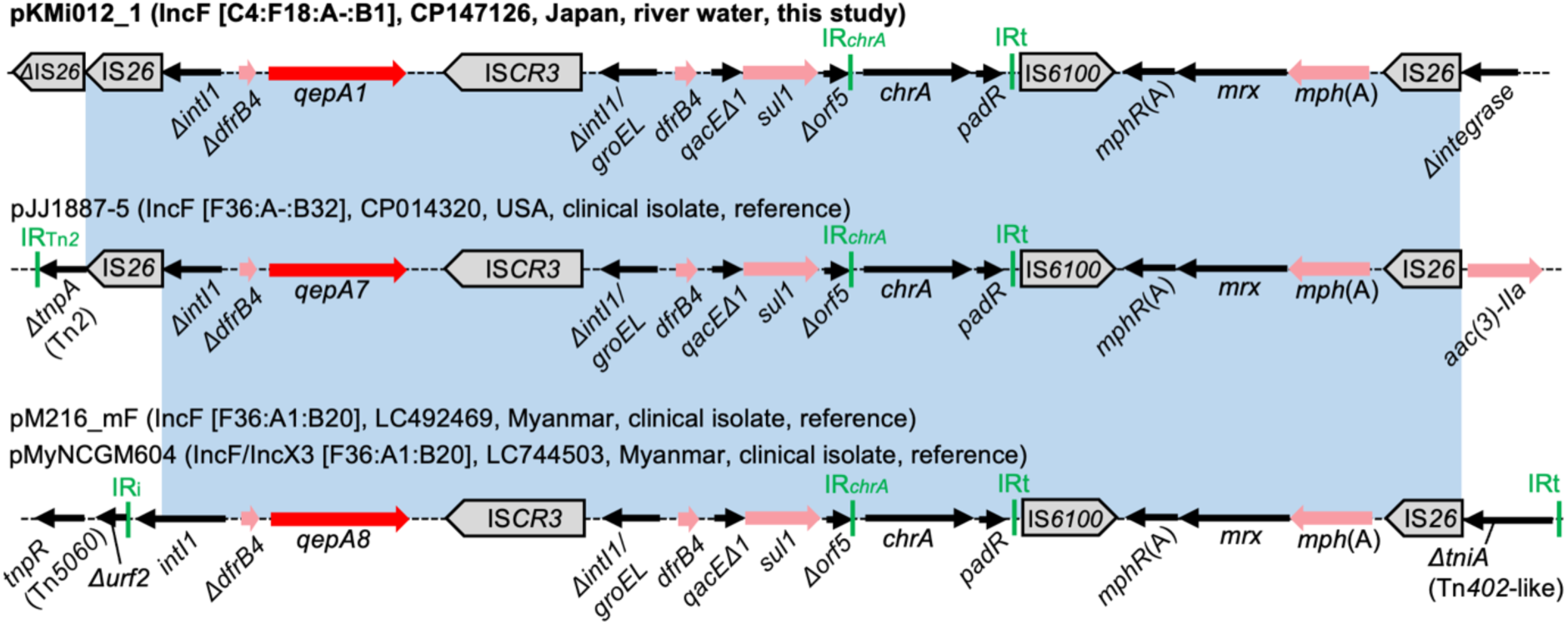
Genetic contexts of *qepA1*. Genes and elements are shown as in earlier figures. The blue shaded regions indicate nucleotide sequences with >99% identity. The contexts detected in three plasmids from clinical isolates are shown for reference purposes. These clinical plasmids carry *qepA7* or *qepA8* instead of *qepA1*. The *ΔintI1/groEL-dfrB4-qacEΔ1-sul1-Δorf5* region, the chromate resistance (*chrA*) region, and the macrolide resistance (*mph*(A)) region are present downstream of *qepA* in all four plasmids.

### 3.13. Study limitations

This study has some limitations. First, a limited number of complete genomes (n = 20) were determined in this study. We also included RefSeq genomes in our analysis to extend our dataset, which might partially mitigate this limitation. Second, there are potential biases in the analyzed isolates. Even though we supplemented our dataset with RefSeq genomes, most RefSeq genomes were from Japan and Switzerland. This seems to reflect the situation where long-read sequencing is still not widely used to study antibiotic resistance in environmental bacteria. Another potential bias lies in the isolation procedure. For example, our nine wastewater isolates and some RefSeq isolates were obtained using selective media for ESBL-producers. This could have influenced the relative abundance of certain genetic contexts (e.g., *qnrS1* genetic context types B–D, which link *bla*_CTX-M-15_ with *qnrS1*). Third, the numbers of complete genomes were small for *qnrA1* (n = 1), *qnrB7* (n = 2), *qnrB19* (n = 2), and *qepA* (n = 1). This might reflect the scarcity of these genes in environmental *E. coli*. However, we also included genomes of clinical isolates in our analysis, which allowed comparative genomic analysis for these genomes. Notably, and while not a limitation of our own study, we unexpectedly found that even though the sequences of these clinical isolates were previously reported, the genetic contexts of TMQR genes were not analyzed in many of the original studies and thus were elucidated for the first time in the present study.

## 4. CONCLUSIONS

Here, we performed in-depth analysis of the genetic contexts of TMQR determinants in environmental *E. coli*. To our knowledge, this is the most comprehensive study analyzing the genetic contexts of TMQR determinants in environmental bacteria. The genetic contexts described in this study included those closely related to the contexts found in clinical isolates. We also detected novel contexts in our genomes (e.g., TIME-COMP containing *qnrD1*) and previously uncharacterized contexts in RefSeq genomes (e.g., a small cointegrate plasmid carrying *qnrB19*). Some TMQR genes were frequently linked to other AMR genes in the described contexts, and this information will contribute to our understanding of how multidrug resistance emerges and spreads in bacteria. We also found multiple examples of TMQR determinants located on the chromosome. Chromosomal location of TMQR genes may contribute to maintenance of these genes, but further studies are needed to confirm this. Overall, this study provides valuable insights into mobile genetic elements associated with TMQR determinants and highlights the importance of genomic surveillance of antimicrobial resistant bacteria in the environment.

## CRediT authorship contribution statement

Conceptualization: R.G.; methodology: R.G.; validation: R.G.; formal analysis: R.G.; investigation: R.G. and F.A; resources: R.G. and F.A; data curation: R.G.; visualization: R.G.; funding acquisition: R.G.; writing – original draft: R.G.; writing – review & editing: R.G. and F.A.

## Acknowledgments

This work was supported by JSPS KAKENHI (grant number JP22K18038), the Kurita Water and Environment Foundation (22B006), the Sumitomo Foundation (2230113), and the Environment Research and Technology Development Fund (JPMEERF20235R01) of the Environmental Restoration and Conservation Agency, provided by the Ministry of the Environment of Japan. Computations were partially performed on the NIG supercomputer at ROIS National Institute of Genetics. We acknowledge the NGS core facility of the Genome Information Research Center at the Research Institute for Microbial Diseases of Osaka University for the support in DNA sequencing. We also thank Isao Nakamoto and Yasufumi Matsumura for their support in antimicrobial susceptibility testing.

