## Supplementary Materials and methods and Supplementary Figures for "Quinolone resistance genes *qnr*, *aac(6′)-Ib-cr*, *oqxAB*, and *qepA* in environmental *Escherichia coli*: insights into their genetic contexts from comparative genomics"

#### Genome sequencing and assembly

Twenty-four *E. coli* isolates, including those from our previous collections and newly obtained ones, were subjected to short- and long-read sequencing in the present study as described below (note that short reads were obtained for some of the isolates in our previous studies but we obtained short reads again in this work because the short reads in our previous studies had low/uneven sequencing depth).

DNA was extracted from each isolate using a DNeasy PowerSoil Pro Kit (Qiagen, Hilden, Germany). Short-read sequencing libraries were prepared using an Illumina DNA PCR-Free Prep kit (Illumina, San Diego, CA). The libraries were sequenced on a NovaSeq 6000 platform (Illumina), generating paired-end reads of 151 bp each. Long-read sequencing libraries were prepared using the SQK-LSK109 kit (Oxford Nanopore Technologies, Oxford, UK) and sequenced on the MinION with FLO-MIN106 (R9.4.1) flow cells.

Short reads were trimmed using fastp (v0.23.2),[1] and long reads were filtered using Filtlong (v0.2.0, https://github.com/rrwick/Filtlong). Hybrid assembly was performed with a long-read-first approach, i.e., long-read assembly using Flye (v2.9.1-b1780)[2] followed by long-read polishing with Medaka (v1.8.0, https://github.com/nanoporetech/medaka) and short-read polishing with Polypolish (v0.5.0).[3] Hybrid assembly was also performed with a short-read-first approach using Unicycler (v0.5.0).[4] If the long-read-first approach generated a complete genome, the polished Flye assembly was chosen as a final assembly. We note that long-read-first assemblies sometimes lack small plasmids, so we recovered small plasmids that were not present in the Flye assembly but were present in the Unicycler assembly.[5] If the long-read-first approach did not generate a complete genome but the short-read-first approach did, the Unicycler assembly was chosen as a final assembly.

If the above approaches failed to generate a complete genome, we performed long-read sequencing again using DNA prepared from the same bacterial pellet but with a different extraction method, which employs enzymatic digestion followed by proteinase K digestion and AMPure XP (Beckman Coulter, Inc. Brea, CA, USA) bead purification, aiming to reduce DNA shearing and to obtain longer reads. The obtained long reads with less fragmentation were subjected to hybrid assembly as described above.

### Supplementary Figures

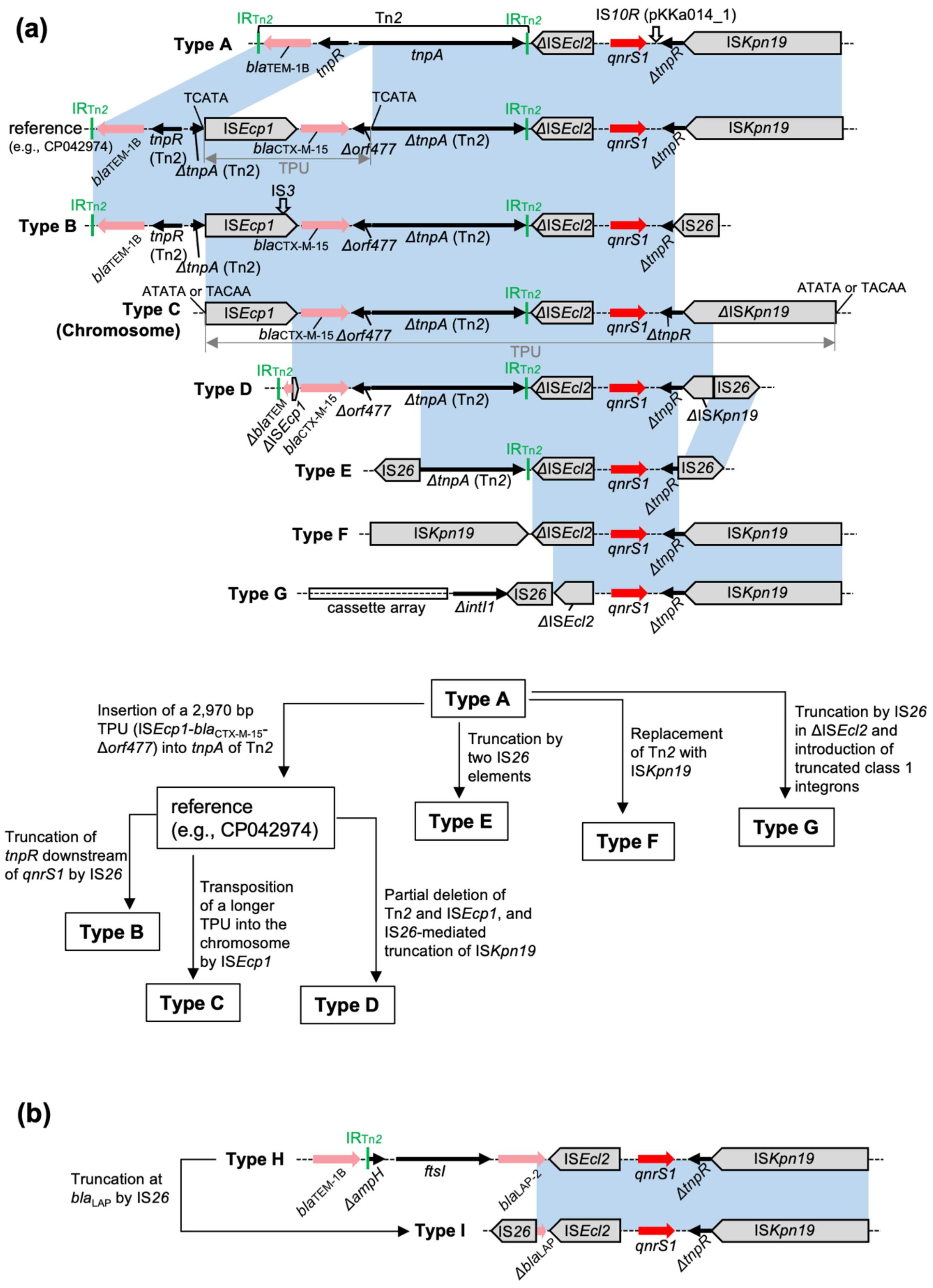

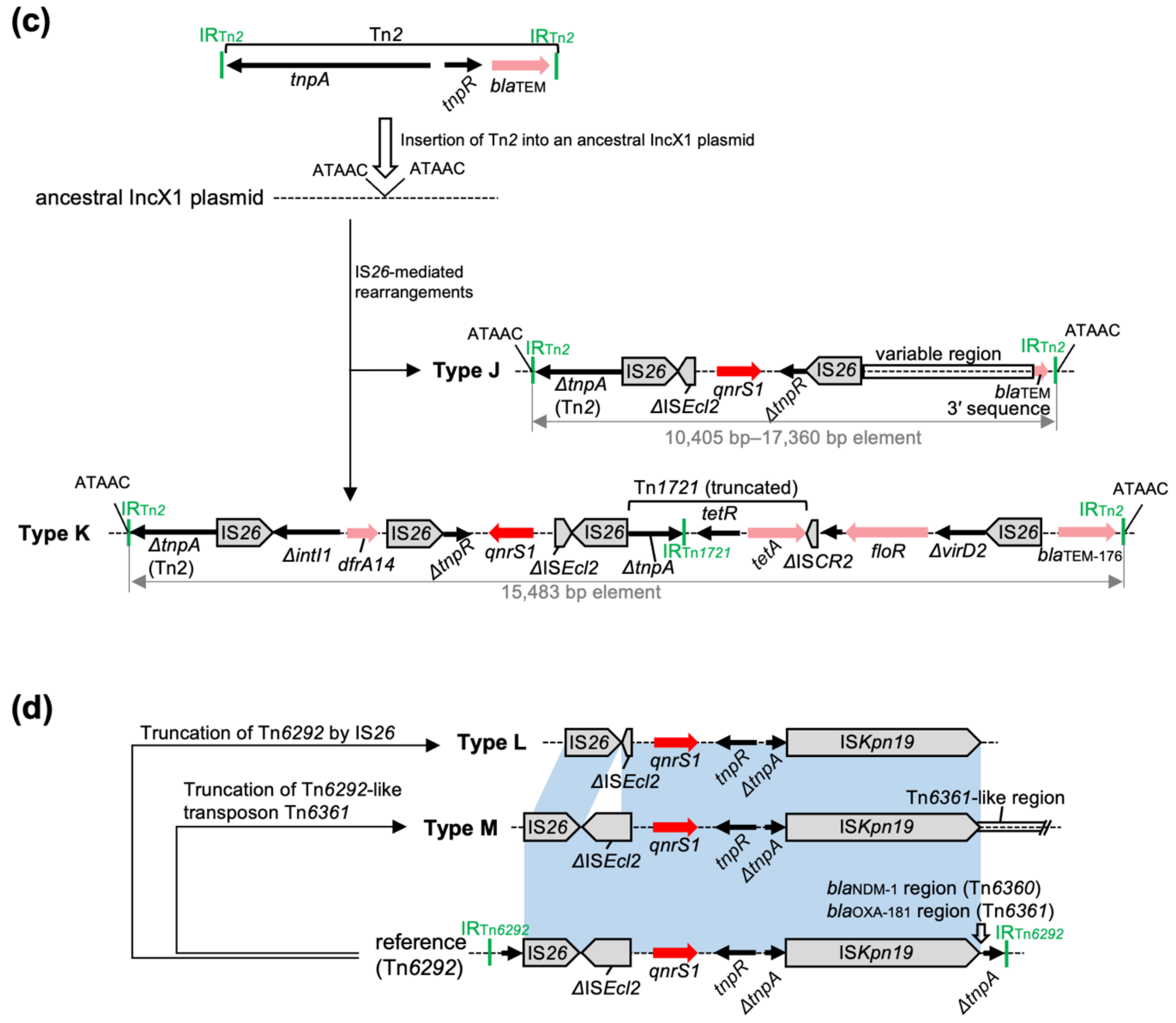


**Figure S1**. Possible evolutionary pathways of *qnrS1* genetic contexts. The contexts in **Figure 4** are shown with reference sequences. (a) The contexts related to type A. Type A is the context detected in the original *qnrS1*-harboring plasmid pAH0376 (AB187515) from a clinical *Shigella flexneri* isolate (note that the available pAH0376 sequence ends at *tnpR* downstream of *qnrS1* and the presence of IS*Kpn19* in the plasmid is unknown).[6] Insertion of a 2,970 bp TPU (IS*Ecp1*-*bla*$_{\text{CTX-M-15}}$-*Δorf477*) into *tnpA* of Tn*2* in type A, which is a context prevalent in genomes deposited in GenBank (e.g., CP042974) but not detected in this study, seems to have been followed by (i) truncation of *tnpR* downstream of *qnrS1* by IS*26* (type B), (ii)

transposition of a 11,384 bp TPU, including the original TPU (IS*Ecp1*-*bla*$_{\text{CTX-M-15}}$-*Δorf477*) and the *qnrS1* region, into the chromosome by IS*Ecp1* (type C), or (iii) partial deletion of Tn*2* and IS*Ecp1* and IS*26*-mediated truncation of IS*Kpn19* (type D). Truncation of type A by two IS*26* elements might have led to generation of type E. In type F, Tn*2* was replaced with IS*Kpn19*. In type G, IS*Ecl2* is truncated by IS*26*, and class 1 integrons with different cassette arrays are present upstream of *qnrS1*. (b) The contexts related to type H. The type H context was detected in multiple *Enterobacteriaceae* plasmids by blastn analysis, and truncation of type H by IS*26* at *bla*$_{\text{LAP-2}}$ seems to have given rise to type I. (c) The contexts found on three IncX1 plasmids. Type J and type K contexts are inserted in the same position in the IncX1 plasmids and flanked by 5-bp direct repeats of ATAAC. Insertion of Tn*2* in the ancestral IncX1 plasmid followed by IS*26*-mediated rearrangements, including insertion of the region containing *qnrS1*, can explain these contexts. In **Figure 4**, the type K context was flipped for the alignment of *qnrS1* regions. (d) The contexts related to Tn*6292*. Type L and Type M seem to have been derived from the *qnrS1*-bearing transposon Tn*6292* (or Tn*6292*-like transposons). Tn*6292* itself was not detected in the present study, though blastn analysis identified that intact Tn*6292* is almost restricted to *Enterobacteriaceae* plasmids from China.
